# Building an atlas of transposable elements reveals the extensive roles of young SINE in gene regulation, genetic diversity, and complex traits in pigs

**DOI:** 10.1101/2022.02.07.479475

**Authors:** Pengju Zhao, Lihong Gu, Yahui Gao, Zhangyuan Pan, Lei Liu, Xingzheng Li, Huaijun Zhou, Dongyou Yu, Xinyan Han, Lichun Qian, George E. Liu, Lingzhao Fang, Zhengguang Wang

## Abstract

Transposable elements (TEs) are an extensive source of genetic polymorphisms and play an indispensable role in chromatin architecture, transcriptional regulatory networks, and genomic evolution. The pig is an important source of animal protein and serves as a biomedical model for humans, yet the functional role of TEs in pigs and their contributions to complex traits are largely unknown. Here, we built a comprehensive catalog of TEs (n = 3,087,929) in pigs by a newly developed pipeline. Through integrating multi-omics data from 21 tissues, we found that SINEs with different ages were significantly associated with genomic regions with distinct functions across tissues. The majority of young SINEs were predominantly silenced by histone modifications, DNA methylation, and decreased accessibility. However, the expression of transcripts that were derived from the remaining active young SINEs exhibited strong tissue specificity through cross-examining 3,570 RNA-seq from 79 tissues and cell types. Furthermore, we detected 211,067 polymorphic SINEs (polySINEs) in 374 individuals genome-wide and found that they clearly recapitulated known patterns of population admixture in pigs. Out of them, 340 population-specific polySINEs were associated with local adaptation. Mapping these polySINEs to genome-wide associations of 97 complex traits in pigs, we found 54 candidate genes (e.g., *ANK2* and *VRTN*) that might be mediated by TEs. Our findings highlight the important roles of young SINEs in functional genomics and provide a supplement for genotype-to-phenotype associations and modern breeding in pigs.

## Introduction

Transposable elements (TEs) or common repeats are ubiquitous sequences that can copy and insert themselves throughout the eukaryotic and prokaryotic genome^1-7^. The movement of TEs is often accompanied by an increase in their abundance, comprising a large fraction of genomic sequence^8-9^. According to the mechanism of transposition, TEs can be generally classified into 1) RNA-mediated class I elements (retrotransposons), including long terminal repeats (LTRs), long interspersed nuclear elements (LINEs), and short interspersed nuclear elements (SINEs); and 2) RNA-independent class II elements (DNA transposons)^10^. TE classes could be further divided into distinct families or subfamilies in terms of their age (active period) and DNA sequence characteristics.

At the predominant view of the 1960s-1990s, TEs were described as “selfish” or “junk” DNA^11^. Thanks to the availability of whole-genome sequences of various species and the ongoing development of bioinformatics tools^12-15^, our knowledge of TEs has progressed at a fast pace. TEs are known to play an essential role in shaping genomic sequences and contributing to the diversity in genome size and chromosome structure^1,16-17^. Most of TEs, in fact, are fixed, inactive, and not randomly distributed in the genome^18-19^. However, several TE families are still actively transposing and serving as a major source of genetic polymorphisms between individuals, such as the Alu, L1, and SVA TE families in the human genome^20^. It is evident in many species (e.g., human, rice and bird) the impacts of active TEs on genome evolution are wide-ranged, including admixture, adaption, footprints of selection, and population structure^21-24^. For example, the polymorphic TEs (polyTEs) detected in the 1000 Genomes Project, consisting of 16,192 loci in 2,504 individuals across 26 human populations, successfully recapitulated the human evolution and captured the sign for positive selection on recent human TE insertions^20,25-26^.

In addition to their direct influences on DNA sequence, there is also emerging evidence that TEs have important functional contributions to gene regulatory networks and epigenome variation. For instance, TEs can directly affect gene transcriptional structure by provoking various forms of alternative splicing, including exonization, exon skipping, and intron retention (3’S and 5’S), to generate novel protein-coding sequences or premature ends^27-30^. TEs can disrupt the existing *cis*-regulatory elements, e.g., promoter, enhancer, and insulator, or provide novel ones^31-34^. They can also serve as a rich source of non-coding RNAs, including LncRNAs, circRNA, small RNA, and microRNA targets^35-38^. Moreover, the silence of TEs has a close connection with epigenetic regulatory mechanisms, such as DNA methylation, piRNA, histone modifications, and RNA interference (RNAi)^18,39-42^. These epigenomic landscapes, together with the TE landscapes, vary from breed to breed in plants, e.g., angiosperms^43^. Importantly, it has been reported that the complex interactions between TEs and epigenetic elements could allow a rapid phenotypic adaptation to environmental changes^40,42,44^.

Pig (*Sus scrofa*), one of the earliest domestic animals, is estimated to be domesticated approximately 10,000 years ago in Asia and Europe independently^45^. It serves as an indispensable source of animal protein and an important biomedical model for humans^46-47^. Currently, a total of 22 pig assemblies are publicly available in NCBI^48-52^, accompanied by the availability of massive high-throughput whole-genome sequences (WGS), providing researchers with ideal materials to boost the current development of genomic research in pigs. However, the study of TEs in the pig genome is still in its infancy. A few previous studies mainly focused on its diversity and distribution^48-52^, yet the functional and evolutionary significance of TEs is largely overlooked in pigs. In our recent study^53^, we identified novel introgressions in Eurasian boars from Asian and European pig populations using the SINE (PRE-1 subfamily) polymorphisms, suggesting that a part of TEs are still active in the current pig genome. Nevertheless, these studies are far from sufficient to comprehensively understand the important roles of TEs in pigs.

In this study, we built the most comprehensive and high-quality atlas of TEs so far in pigs using the newly built pipelines and further defined the SINE families into four categories based on their ages. We then systematically explored the functional aspects of these SINE categories by combining large-scale multi-omics data from 21 tissues, including three-dimensional (3D) chromatin architecture, chromatin accessibility, histone modifications, transcription factor binding sites (TFBS), and DNA methylation. We estimated the contribution of active SINEs to tissue-specific gene expression by cross-examining 3,570 published RNA-seq samples from 52 tissues and 27 cell types. Furthermore, we built a comprehensive atlas of polymorphic SINEs using 374 publicly available WGS data to investigate the roles of young SINEs in pig population admixture and local adaptation. The TE-mediated adaptation has been found in functional regions, such as the almost fixed polySINEs observed in laboratory-inbred Bama Xiang pigs at the upstream region of the *LEP* gene. Finally, by mapping these polySINEs to 4,072 loci associated with 97 complex traits in pigs, we proposed 54 candidate genes that might regulate complex traits through TEs.

## Results

### Composition of young SINE families in the pig genome

To detect TEs as thoroughly as possible, we developed the **Pig TE D**etection and **C**lassification (PigTEDC) pipeline (**Fig. 1a**, Supplementary Information) and applied it to the high-quality pig genome (*Sus scrofa* 11.1). The PigTEDC pipeline employed a combination of similarity-, structure-, and *de novo*-based methods. Based on the existing TE repository (RepBase update and Dfam 2.0 databases), we further classified all potential pig TEs into classes/superfamilies and families and derived their consensus sequences.

**Figure 1.**
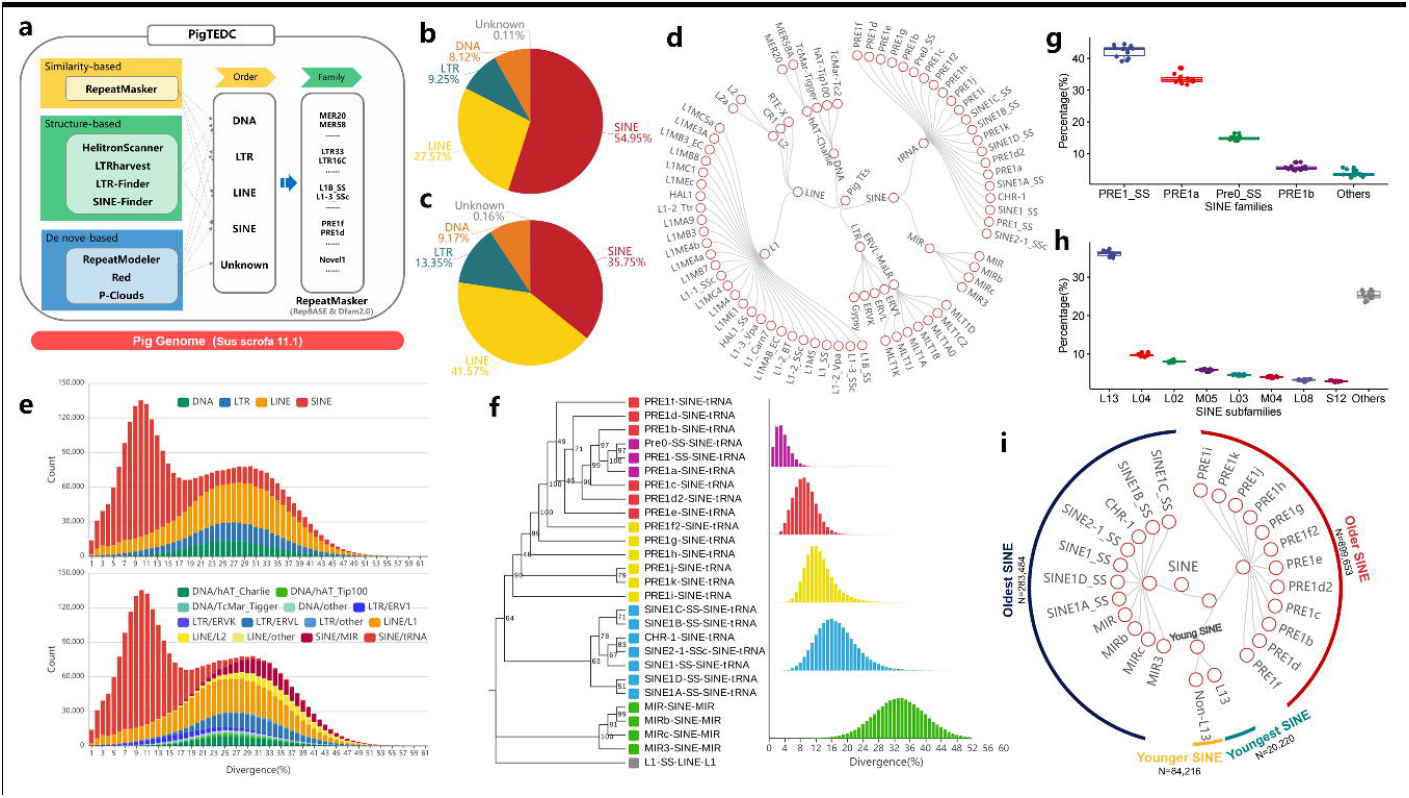
Transposable elements annotation and SINE classification in the pig genome. **a** A schematic of the Pig Transposable Element Detection and Classification (PigTEDC) pipeline. It is composed of three TE detection approaches, using the similarity-, structure-, and *de novo*-based algorithms. **b** The proportion of TEs from different superfamilies in the count. **c** The proportion of TEs from different superfamilies in length. **d** Classification of pig TE superfamilies and families (≥ 3,000 TE copies in each family). **e** Sequence divergence distribution for TE superfamilies (upper panel) and families (bottom panel) in the pig genome. Sequence divergence distributions are plotted in bins of 0.01 increments. **f** Phylogenetic tree and sequence divergence distribution for SINE families in the pig genome. On the right panel, the x-axis represents the divergence, and the y-axis represents the counts of the SINE families. **g** Boxplots displays the proportion of genomic SVs formed by different SINE families. **h** Boxplots displays the proportion of genomic SVs formed by different SINE new subfamilies. See Table S1 for their definitions. **i** Classification of pig SINE families based on their ages.

Excluding the nested TEs, we detected a total of 3,087,929 TEs, occupying 37.9% (947 MB) of the total pig genome. Two-thirds of those TEs were assigned to a specific family. Similar to previous studies^48-52^, the most abundant TEs in the pig genome were retrotransposons (∼90% of TEs), including LTR (9.25%), LINE (27.57%), and SINE (54.95%), whereas DNA transposons only accounted for 8.12% of TEs (**Fig. 1b**). Although SINE had the largest count, its genome coverage was still less than LINE’s (**Fig. 1c**). Out of 532 families of TE, 65 (more than 3,000 TE copies in each family) consisted of 84.6% classified TEs (**Fig. 1d**), particularly for PRE1f in SINE/tRNA (170,511 copies), MIR in SINE/MIR (45,927 copies), L1B-SSc in LINE/L1 (35,819 copies), and MLT1D in LTR/ERVL-MaLR (7,866 copies).

We performed the divergence analysis for all classified TEs using RepeatMasker after the CpG content correction. The stacking plots show the divergence distribution for either superfamilies or families (**Fig. 1e**). Similar to the divergence distribution of TEs in the human genome^54^, we observed two bursts at 10% and 30% in pig TE amplification that were estimated to occur 20 and 60 million years ago (Mya), assuming a substitution rate of 5×10^−9^ substitutions/site per year^55^. Obviously, the amplification of most TE families occurred 70-50 Mya (divergence at 30 ± 5%), consistent with that the Paleocene Epoch (65-54 Mya) opened up vast ecological niches for surviving mammals, birds, reptiles, and marine animals^56^. The latest obvious burst of TEs was mainly concerned with SINE/tRNA, LINE/L1, and LTR/ERV1 families, of which SINE/tRNA was still most active in the modern pig genome^57-59^. Further exploring the ages of highly homologous subfamilies in SINE classes (**Fig. 1f** with an average divergence of 4%, labelled in purple), we found that 3 out of 26 SINE families (PRE1-SS, PRE0-SS, and PRE1a) were recently active and had been proved to be polymorphic within pig breeds in a previous study^53^, and thus can be viewed as the young SINE families.

We next focused on these young SINE families (PRE1-SS, PRE0-SS, and PRE1a) to further reclassify them into subfamilies at a high resolution. After processing all of the full-length young SINEs from 14 publicly available pig genomes (Supplementary Information), we retained 978,506 non-redundant young SINEs and created their consensus sequence by multiple sequence alignment (**Fig. S1-S2**). Subsequently, a minimum spanning (MS) tree analysis recategorized them into 90 new SINE subfamilies, including 17 large (size > 10000), 12 medium (size ≥ 3000 and ≤ 10,000), and 61 small (size < 3000) subfamilies, with *P*-values for subfamily partition ranging from 1×10^−53186^ to 1×10^−4^ (**Fig. S3-S4**, Table S1).

When we compared locations of the polymorphic young SINEs with those of medium-length structure variants (SVs, range from 200 bp to 300 bp) detected from the 14 assemblies as compared to the pig reference assembly (Supplementary Information), we found that polymorphic SINEs mainly belonged to PRE1-SS, PRE1a, and PRE0-SS families (**Fig. 1g**) (on average 90.75% of the medium-length SVs), especially for the L13 subfamily (**Fig. 1h**) (on average 36.15% of the medium-length SVs). This suggests that only a specific set of recently active SINE subfamilies was predominant in contributing to SVs (around 250 bp length) among diverse breeds during the recent evolution in pigs. We therefore further classified all SINEs into four categories of youngest (L13 subfamilies), younger (young SINE families except for L13 subfamilies, non-L13), older (non-young PRE families), and oldest (non-PRE families; ancient families) SINEs. (**Fig. 1i, Fig. S5**).

### Widespread roles of young SINEs in gene regulatory networks

Previous studies proposed that TEs might be co-opted into regulatory sequences of genes *via* diverse epigenetic mechanisms^60-63^. To test this, we explored the impact of SINE subfamilies on the genome features regarding three-dimensional (3D) chromatin architecture (FR-AgENCODE^64^), chromatin accessibility, histone modifications, transcription factor binding sites (TFBS), and DNA methylation (**Fig. 2a**).

**Figure 2.**
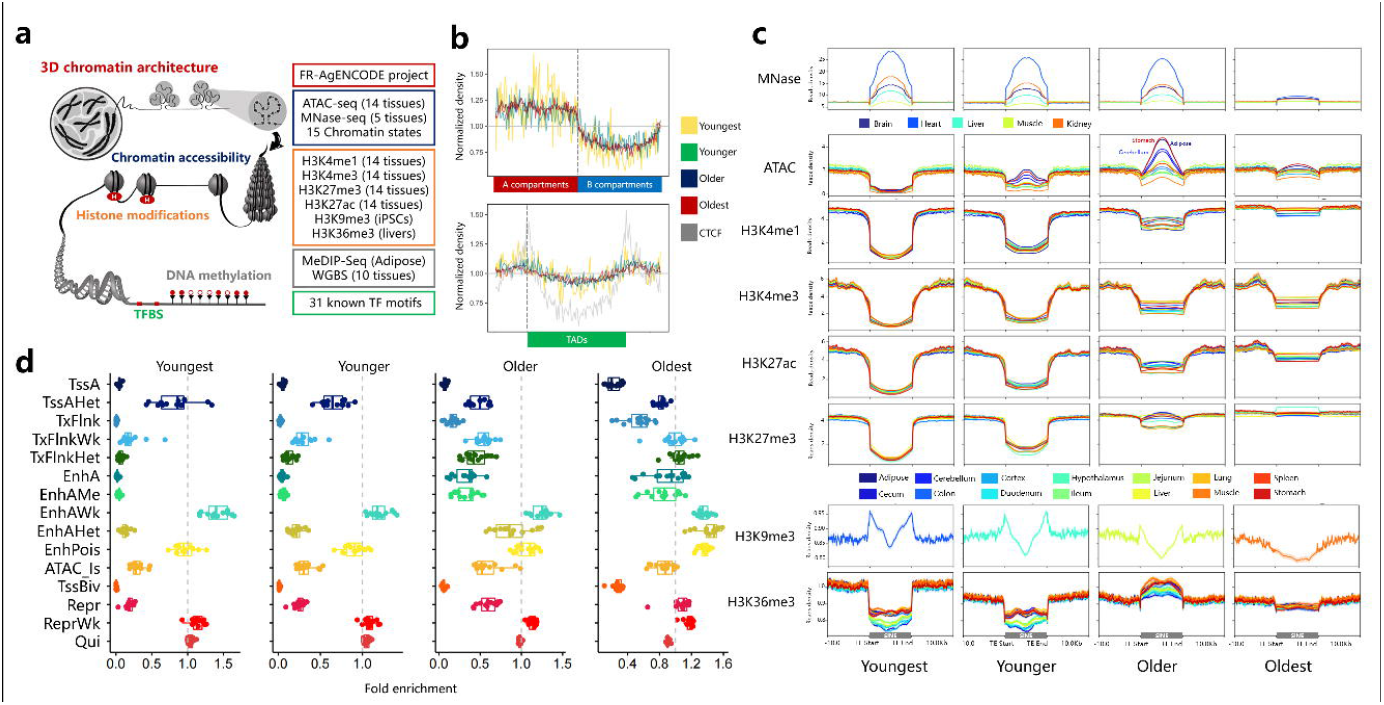
Distribution of SINE on pig genome and functional regions. **a** The five types of genomic features used in this study included 3D chromatin architecture, chromatin accessibility, histone modifications, DNA methylation, and transcription factor binding sites. **b** The distribution of SINE families between 3D chromatin architectures (Compartments A vs. B), as well as near topologically associating domain (TAD). **c** The reads density distributions of chromatin accessibility and histone modifications near transcripts across four different SINE groups. **d** Boxplots displays the enrichment of four SINE groups in 15 distinct chromatin states across 14 tissues.

We observed a highly significant enrichment of all SINEs in the A compartments (active), whereas a depletion in the B compartments (inactive) (Wilcoxon test, *P-*values < 10^−16^, **Fig. 2b, Fig. S6**). After the further separation of A/B compartments into topologically associating domains (TADs), as expected, we observed CTCF binding sites were significantly enriched in the boundary regions of TADs. SINEs showed a similar but slightly weaker trend of enrichment, while the young SINEs showed a higher but more variable enrichment than the old ones (**Fig. 2b, Fig. S7**).

Next, we explored the distribution of SINE families on the chromatin accessibility and nucleosome positioning near transcripts using the published ATAC-seq (14 tissues) and MNase-seq (five tissues) datasets, respectively^65-66^. TE enrichment in open and closed chromatin exhibited strong age-specific patterns, mainly reflecting that the youngest and younger SINE families were significantly depleted from open chromatin regions and enriched near the nucleosome (**Fig. 2c**). Whereas the older SINE families showed relatively high enrichment for open chromatin, particularly in the stomach, followed by adipose and cerebellum.

We further explored the relationship of SINE families with four active epigenetic marks (H3K4me1 - primed enhancers, H3K4me3 - enriched in transcriptionally active promoters, H3K27ac - distinguishes active enhancers from poised enhancers, and H3K36me3 - actively transcribed gene bodies) and two repressive marks (H3K9me3 - constitutively repressed genes and H3K27me3 - facultatively repressed genes). In **Fig. 2c**, we found that the majority of SINE families were underrepresented in all four active marks, consistent across tissues. Whereas SINE families, particularly for young SINEs, were highly enriched for H3K9me3 but not for H3K27me3. H3K27me3 has development-dependent repressed characteristics, while H3K9me3 indicates permanent repression^67^. In general, compared to older or oldest SINE groups, younger and youngest SINE groups (especially the youngest group) showed a much higher over-/under-representation for all types of histone modifications. In addition, we further investigated the enrichment of SINE families in 15 distinct chromatin states across 14 tissues (**Fig. 2d**, Supplementary Information)^68^. We observed that young SINE groups (youngest and younger) show lower enrichment in most of the functional chromatin states than old SINE groups (older and oldest), and the enrichment degree of SINE in chromatin states was roughly similar across 14 tissues. These enrichment characteristics can be further divided into four distinct enrichment patterns according to the enrichment degree of different SINE groups (**Fig. 3a)**. We observed that the oldest SINE’s highly enriched pattern accounted for the vast majority (80%, 168 of 210 combinations of 14 tissues and 15 states), while the young SINE group showed the three enrichment patterns in remaining combinations (the enlarged inset). In two of them, only the youngest SINE group showed high enrichment for TssAHet (flanking active TSS without ATAC) and EnhAWk (weakly active enhancer). In general, all SINE groups were significantly depleted from active promoters and enhancers, except for weak TSS and enhancers, and the young SINE groups showed a higher depletion than old ones. All these together indicated that young SINEs as new invaders might be silenced by histone modifications and DNA methylation, while the old SINEs might be tolerated by the pig genome.

**Figure 3.**
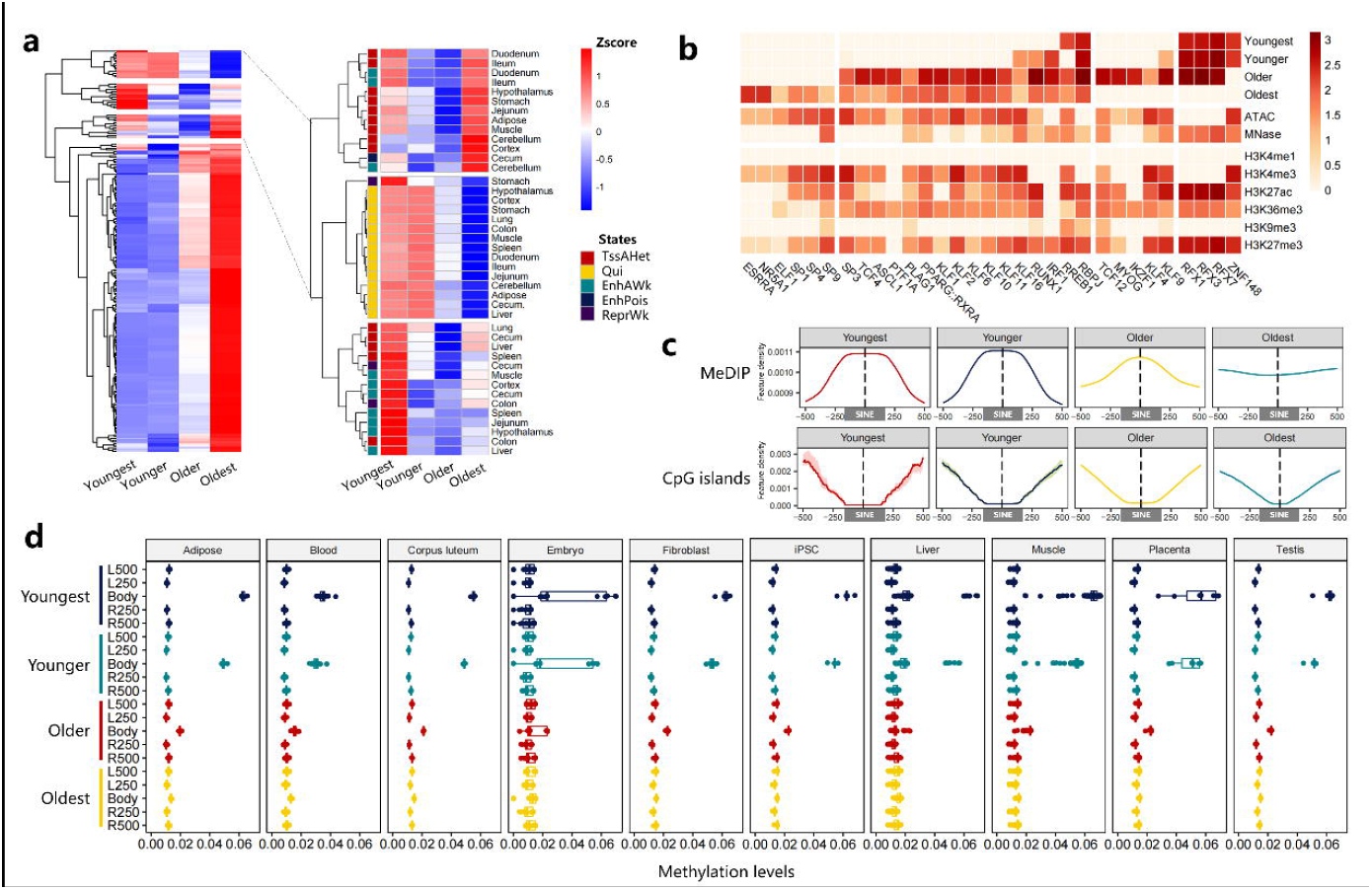
Enrichment of SINE in functional elements and methylation modification. **a** Hierarchical clustering of enrichment patterns in 15 chromatin states for four SINE groups across 14 tissues (left panel). The Heatmap of three distinct enrichment patterns for the high enrichment in the young SINE group (right panel). **b** A heatmap for the enrichment of transcription factor binding motifs in SINE families, chromatin accessibility, and histone modifications. **c** The signal density of MeDIP-seq and CpG island within different SINE families. **d** Boxplots displays the DNA methylation levels on different SINE families. L and R represent the upstream and downstream directions of the SINE body. E.g., L250 represents 0 to 250 bp, and L500 represents 250 to 500 bp window upstream of SINE.

It has been proposed that TEs carrying TFBS repertoire may indirectly contribute to the transcriptional regulation of genes^69-70^. We thus performed the motif enrichment analysis of SINEs to determine their possible contributions (**Fig. 3b**). In total, 31 known TFBS were predicted to have the binding motifs in at least one SINE family, 30 of which (96.8%) were found in old ones and 26 (83.9%) were related to the open chromatin, revealing that the recent exaptation of young SINEs into regulatory regions was relatively rare^71^ and was repressed by less chromatin accessibility. The youngest SINE-specific TFBS related to the *ZNF148* gene has been proven to drive the formation of a muscle phenotype^72^. Besides, three members of the RFX transcription factor family were observed to be amplified in young and older SINEs and involved in the immune, reproductive, and nervous pathways^73^. For example, RFX1 and RFX3 were the candidate major histocompatibility complex (MHC) class II promoter binding proteins that were found to function as a *trans*-activator of the hepatitis B virus enhancer^74-75^.

Given that TEs may play major roles in the regulation of gene expression by shaping the epigenetic modifications^76^, we further assessed the epigenetic states of SINE families in terms of DNA methylation (MeDIP), the density of CG (CpG) sequence contexts, and AT:GC content (CpG islands) (**Fig. 3c**; Supplementary Information). The results showed that almost all SINE families exhibited a significant depletion of CpG islands, and the young SINE families were more highly methylated than the old ones. Similarly, we observed that the average CG methylation levels within SINE bodies, especially for the young SINEs, were significantly higher than their flanking regions across 10 different tissues, which corresponded to the enrichment of SINE families in H3K9me3 (**Fig. 3d**). A previous study revealed that Piwi-interacting RNAs (piRNAs) played a major role in TE silencing via the ping-pong cycle in pig germline^77^. We further distinguished three classes of small non-coding RNAs to test the relationship between piRNA density and SINE families (**Fig. S8**, Supplementary Information). In contrast to siRNA and miRNA, piRNAs were significantly enriched for SINE-related sequences or sequence flanks, and there was a significant negative correlation between piRNA density and the age of SINE subfamily. Our result was in line with the findings in humans that young SINE families were more likely to be targeted by piRNAs^78^, which can be regarded as the major reason leading to the high methylation levels of young SINE families, as we observed above.

### Young SINE-derived transcriptome profiling in pigs

In addition to their interactions with epigenetic modifications, TEs can directly modify the transcription of host genes by remodeling new alternative splice events^79-80^. To test this, we analyzed the PacBio long-read isoform sequences (Iso-Seq) from 38 pig tissues^81-82^ using a uniform pipeline to detect the transcripts of SINE-derived exonization and alternative splice sites. We estimated the contribution of TEs to gene expression across 52 tissues and 27 cell types by analyzing 3,570 published RNA-seq samples (**Fig. 4a**, Table S2) from the EBI database (https://www.ebi.ac.uk/).

**Figure 4.**
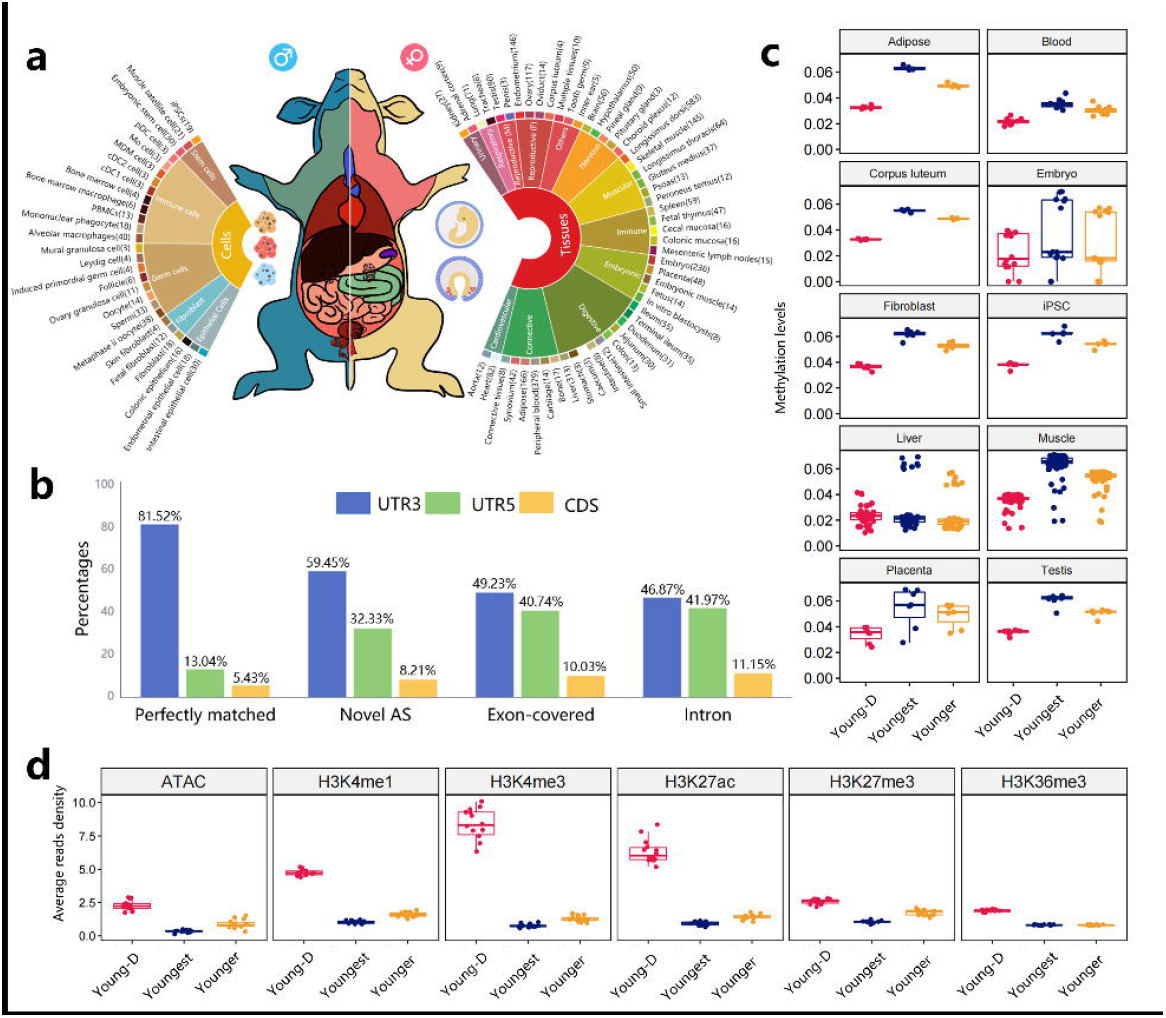
Young SINE-derived transcriptome landscape. **a** Overview of RNA-seq libraries in 3570 samples across 52 tissues and 27 types of cells. **b** The bar plot indicates the proportion of functional regions affected by SINE across four different categories of SINE-derived transcripts. **c** Boxplots display CG methylation levels on young SINE families. Young-D group represents the SINE families that derived the young SINE-derived transcripts. The Younger and Youngest groups represent all the younger and youngest SINEs in the entire genome, respectively. **d** Boxplots displays the reads density of chromatin accessibility and histone modifications on Young-D, Younger, and Youngest groups.

After processing raw data by LoRDEC^83^, we obtained 30,331,870 error-corrected Iso-seq reads with a mean length of 2,797 bp, of which 7.48% (2,267,973) was defined as TE-derived transcripts. Importantly, 68.81% (1,560,568) of TE-derived transcripts were recognized to be inserted by nearly full-length SINE (average coverage of 87.76%), indicating that SINE plays an important role in regulating gene expression.

Similar to a previous study^82^, we next classified 337,746 young (younger and youngest) SINE-derived transcripts into four categories by comparing their genomic location with known transcripts in the currently available pig genome annotations (**Fig. S9**). Out of those, 1,028 young SINE-derived transcripts perfectly matched with 517 PCGs and 47 LncRNAs (Table S3), and 62,304 young SINE-derived transcripts potentially offered the novel alternative splice events for 8,103 PCGs and 405 LncRNAs (overlapping with at least one splice junction of a known transcript). The remaining young SINE-derived transcripts with no complete structural similarity with the currently available transcript annotation were classified as either 150,469 exon-covered (exonic overlap without any splice junction on the same or opposite strands) or 130,180 intronic transcripts (felled in a reference intron).

To understand the *cis*-functionality of young SINE-derived transcripts, we used the transcriptome data from 52 tissues and 27 cell types to quantify the abundance (RNA-seq counts) of PCGs and SINE-derived transcripts by Salmon tools^84^. A total of 3,138 protein-coding transcripts (3,112 PCGs) had significant associations with young SINE-derived transcripts at a Bonferroni significance threshold of 1.42×10^−6^ (0.05/35,135). Of those, 84 PCGs showed that the young SINE-derived exonization and their SINE-derived exons were involved in the expression of PCGs (Table S4). Interestingly, we found that the majority of SINE-derived exons (81.52%) were presented in mRNA 3′-untranslated regions (3′-UTRs) (**Fig. 4b**), suggesting that young SINEs can directly insert into the regulatory region to influence gene expression by the mechanism similar to Staufen-mediated decay (SMD)^85^. For example, we found that a full-length PRE0-SS (sus-specific SINE) was inserted in the 3′-UTR of the pig *PDK1* gene, which was in agreement with the previous report that the Alu and B1 (primate-specific and rodent-specific SINEs, respectively) regulated both human and mouse orthologs of *PDK1* by SMD^86^. This provides further supports for the convergent evolution of SINE-directed SMD. In contrast, a higher relative proportion of young SINEs was found in the CDS regions of exon-covered and intronic SINE-derived transcripts (10.03% and 11.15%, respectively) (**Fig. 4b**), suggesting that the selections against the young SINEs might be more relaxed in these two types of SINE-derived transcripts. However, it was consistent that most young SINEs do not directly participate in the protein translation but indirectly influence gene expression by affecting the UTRs of their derived transcripts^87^.

We then retrieved young SINEs involved in SINE-derived transcripts and named them young-D SINEs. We observed that the average CG methylation levels of young-D SINEs that derived transcripts were significantly lower than the entire young SINEs in most tissues (**Fig. 4c**). Similarly, young-D SINEs were more significant enrichment in open chromatin and histone modifications than the entire young SINEs, especially the H3K4me3, providing more evidence that young-D SINEs and their derived transcripts were more likely active and functional across tissues (**Fig. 4d**).

Based on the normalized expression of PCGs by DESeq2^88^, the *t*-SNE plots of unsupervised clustering of 3,570 samples clearly reflected tissue types (**Fig. 5a**). We further performed a co-expression network analysis of 14,403 PCGs using WGCNA R package^89^ to explore the function of young SINE-derived transcripts across this wide range of tissues and cell types (**Fig. S10**). As a result, a total of 13,872 PCGs were grouped into 40 modules with the gene size ranging from 30 to 1,694 (**Fig. S11**, Table S5), and most of the modules showed high tissue specificity and likely play key roles in particular organ systems in pigs (**Fig. S12-S13**). The results were also supported by the gene-to-gene networks of topological clustering, which was performed using the Markov clustering (MCL) algorithm^90^ (**Fig. S14**). Importantly, 17.9% of co-expressed PCGs (2,744) were related to young SINE-derived transcripts and were mainly enriched in modules M38 (Trachea), M7 (Adipose), M18 (Fetal thymus), and M12 (Alveolar macrophages) (Z-score > 1), and the expression of all those modules showed high tissue specificity (**Fig. S15**).

**Figure 5.**
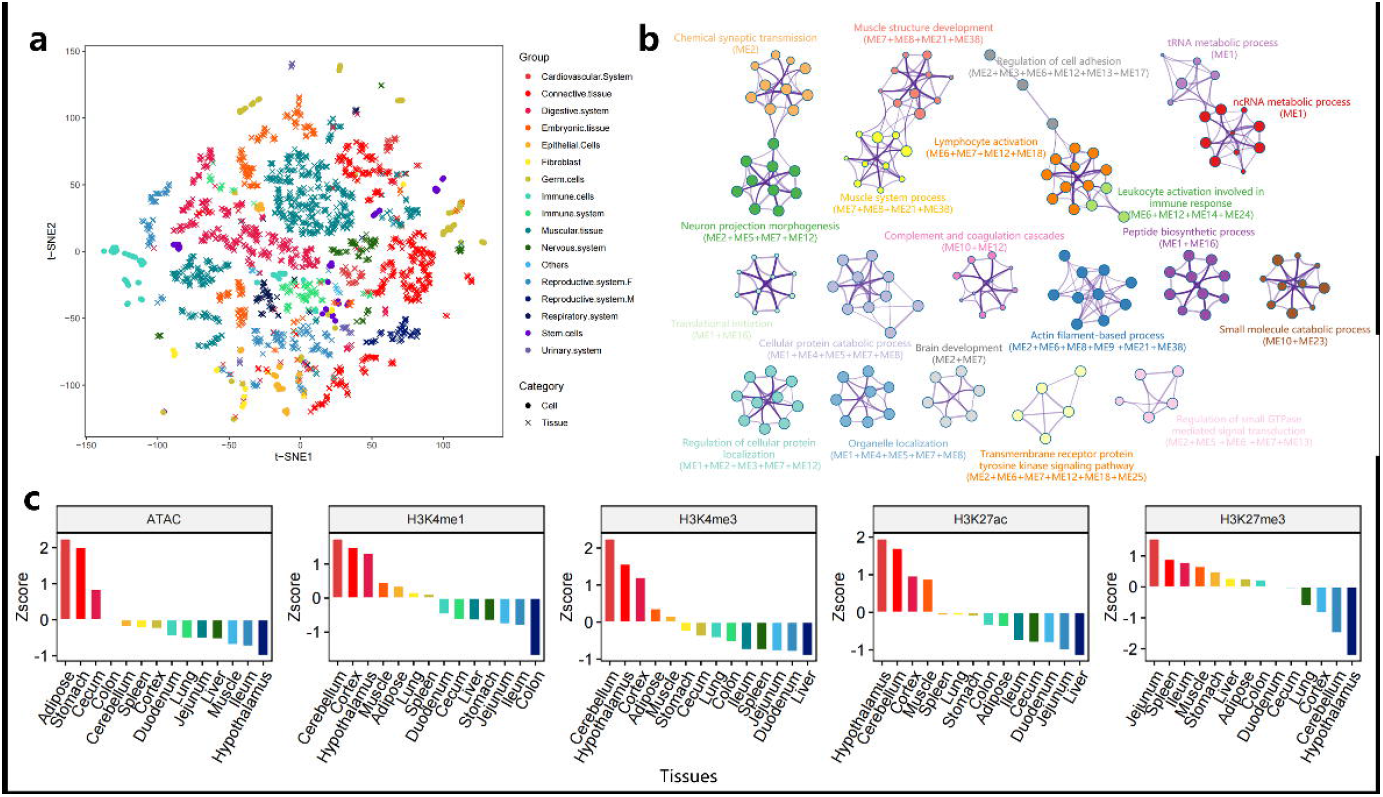
Functional enrichment of young SINE-derived transcripts. **a** The tSNE plots display the expression differentiation among different tissues and cells. **b** Top 20 results of functional enrichment analysis for young SINE-derived genes. **c** The bar plot indicates the enrichment of 248 young-D SINEs in chromatin states and histone modifications across different tissues.

In addition, as shown in **Figure 5b**, these young SINE-related PCGs were significantly enriched in the neural development, cellular metabolism, muscle development, and immune response, which may be responsible for the natural selection and domestication of modern pigs^91-93^. For instance, 186 PCGs were significantly enriched in the chemical synaptic transmission (GO:0007268), brain development (GO:0007420), and neuron projection morphogenesis (GO:0048812), which were mainly in the module M2 with a high expression in brain tissues (**Fig. S16**). Correspondingly, a total of 248 young-D SINEs, whose SINE-derived transcripts had significant associations with these 186 PCGs, were more significantly enriched in the active epigenetic marks (H3K4me1, H3K4me3, and H3K27ac) and depleted from the repressive mark (H3K27me3) at the nervous system (cerebellum, cortex, and hypothalamus) than other tissues, suggesting that young SINEs exhibited strong and concordant tissue specificity in both transcript expression and epigenetic regulation (**Fig. 5c**).

### The roles of young SINEs in population admixture and local adaptation in pigs

TEs produce abundant raw materials for evolution in natural populations, and the burst of TEs was tightly related to significant evolutionary events such as the population admixture and local adaptation^23,94-95^. Young SINE polymorphisms represented the vast majority of TE polymorphisms in the pig genome^48,57,59,96^. Therefore, we developed a comprehensive map of polySINE from WGS of over 300 pigs, representing the majority of Eurasian pig breeds, to explore the roles of young SINEs and their derived genes in pig population admixture and local adaptation.

To investigate whether the sequencing depth and polySINE detection tools had a prominent effect on the detection of polySINEs, we first benchmarked the polySINE detection tools that showed superior performances in previous human projects^13,97^ under different sequencing depths (**Fig. S17-S25**). Based on the results of benchmarking, we customized the **P**ig **TE P**olymorphism (PigTEP) pipeline to maximize its performance in the current pig WGS datasets (**Fig. 6a**). We then applied it to identify polySINEs from 374 individuals with the uniform sequencing depth of 10× (**Fig. S26**, Table S6, average mapped bases: 27.18 GB and average mapping rates: 99.44%). The analyzed 374 individuals from 25 diverse populations (N ≥ 5) were further assigned into 10 major groups, i.e., PYGMY, ISEA, CHD, KOD, AWB, TWB, EUD, EWB, MINI, and COM (**Fig. 6b**).

**Figure 6.**
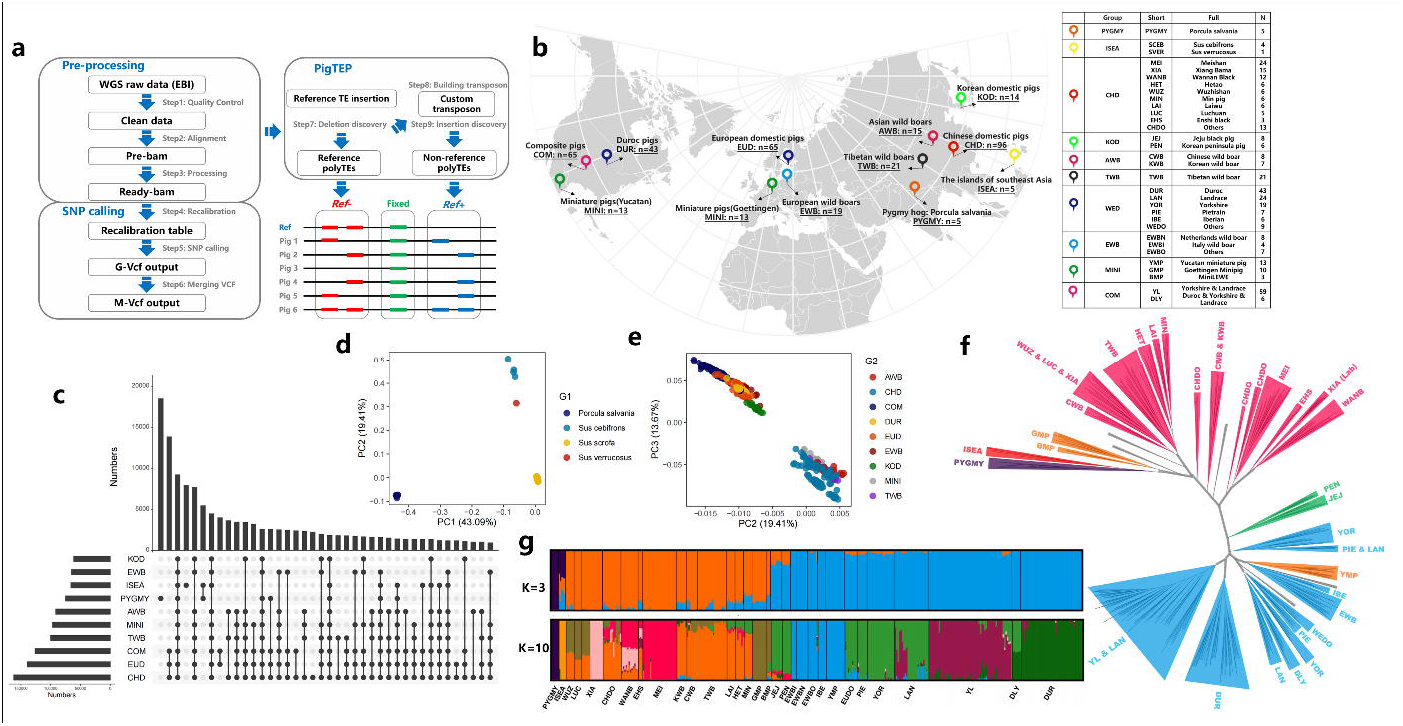
Young SINE-derived genetic diversity of pigs. **a** The Pig TE polymorphism pipeline. The pipeline was constructed to identify both polySINEs and SNPs for each individual simultaneously. **b** Overviews of whole-genome re-sequencings in 374 individuals. **c** Venn plot represents the distribution of polySINEs among different populations. **d** PCA plot displays the genetic relationship based on polySINEs among 374 individuals. **e** PCA plot displays the genetic relationship based on polySINEs among 364 individuals from the modern pigs (*Sus scrofa*). **f** Phylogenetic tree based on polySINEs for 374 individuals. **h** Population structure based on polySINEs for 374 individuals when K was 3 and 10.

In total, we identified 211,067 polySINEs (189,966 *Ref*+ and 21,101 *Ref*-) in pig genome, 49.64% of which were located in non-intergenic regions. As expected, the vast majority of polySINEs were found at a low frequency (64.89% of polySINEs with < 5% minor allele frequencies, MAF) in the whole pig population (**Fig. S27**), but showed variable MAF among groups (**Fig. S28**). We found 85.58% of polySINEs were shared among groups, while only 30,441 polySINEs (PYGMY and ISEA accounted for 60.83% and 26.01%, respectively) were exclusively presented within a single group (**Fig. 6c**).

The principal component analysis (PCA) of samples using polySINE genotypes clearly reflected the four species of the Suidae (**Fig. 6d**). PC1 separated the *Porcula slavania* from *Sus*-species, while PC2 (19.41%) and PC3 (13.67%) showed the genetic separation between Asian and Western breeds (**Fig. 6e**). Interestingly, the Korean domestic pigs (KOD) separated from the Asian breeds and were closer to Western breeds than to Chinese breeds, suggesting the presence of gene flow and introgression from Western pig breeds to Korean domestic pigs and most likely mediated by humans.

The results were further supported by the TE-based phylogenetic tree and genetic admixture (**Fig. 6f and g**; **Fig. S29**), whose evolutionary relationships were basically consistent with previous studies on SNP-based genotypes^45,98^. The comparison of Chinese and European domestic pigs confirmed our previous findings on the TE-based introgression between Northern Chinese domestic pigs and European domestic pigs^57^. Specially, we found that Korean wild boars, unlike the Korean domestic pigs, clustered together with other Asian wild boars instead of European pigs.

To detect polySINEs associated with local adaptation, we calculated the pairwise *Fst*^*i*^ value for each polySINE between cluster i and the remaining clusters to measure its locus-specific divergence in allele frequencies. The polySINEs with extreme *Fst*_*i*_ (top 1%; n =337) were observed in functional regions (exonic, splicing, UTR5, UTR3, and upstream) of 330 PCGs (Table S7). While 77.94% of these PCGs existed in PYGMY (n = 223) and ISEA (n = 42), and the remaining 75 were likely associated with the breed-restricted phenotypes of domestic pigs (*Sus scrofa*) (**Fig. 7a**). For instance, a PRE1 insertion in the promoter of the *IGFBP7* gene, which was associated with tumor suppressor^99^, was widespread in Chinese indigenous breeds rather than in commercial breeds^100^. The upstream (high signals in H3K4me1 and H3K27ac) of *FRZB* was inserted by a population-specific polySINE from Southern Chinese domestic pigs, which was associated with pig growth traits^101^ (**Fig. 7b**).

**Figure 7.**
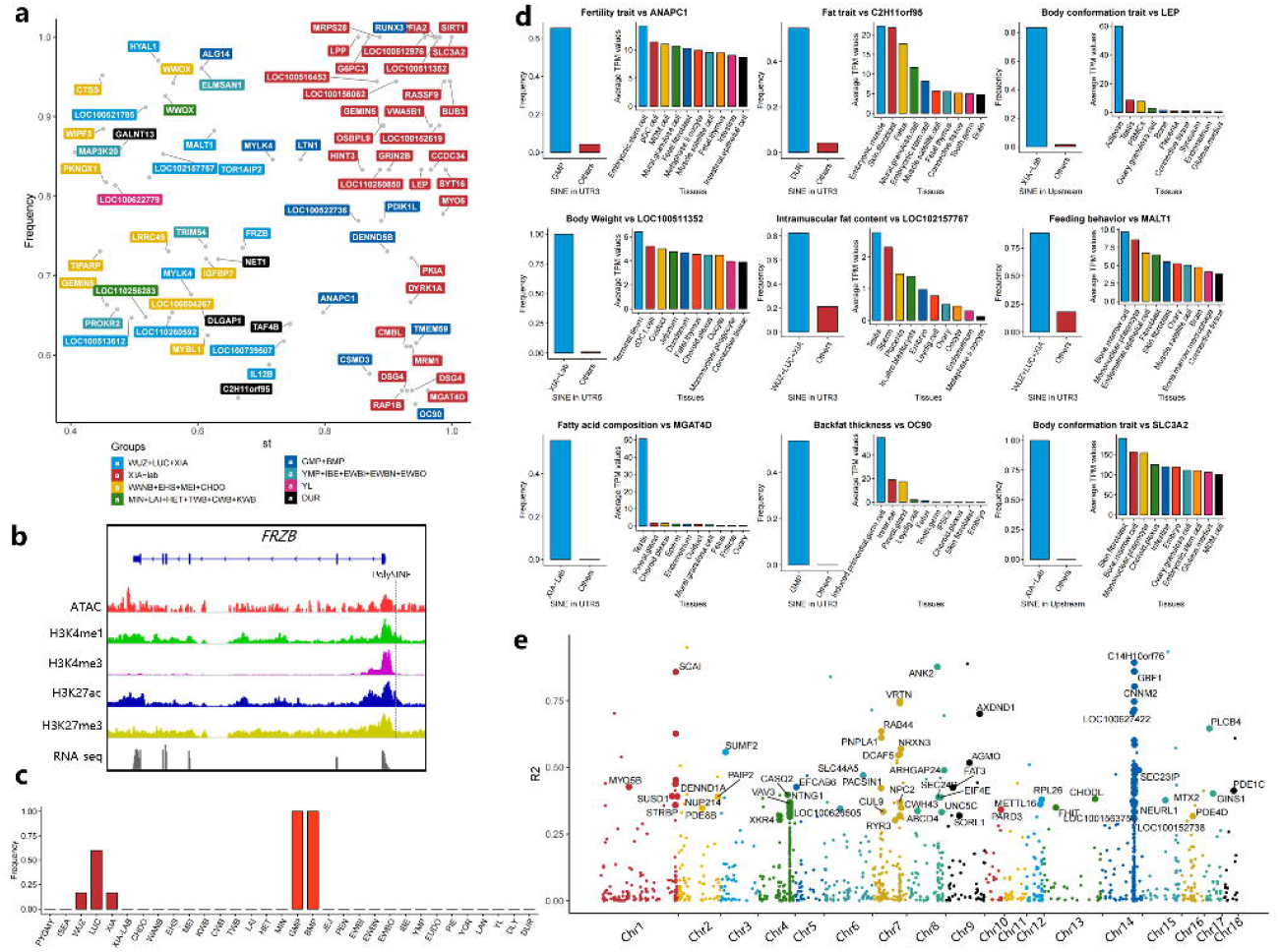
Potential candidate genes for young SINE-derived local adaptation. **a** The scatter diagram displays the 75 PCGs that are possibly associated with local adaptation. The x-axis represents the *Fst*, and the y-axis represents population frequency. **b** Chromatin accessibility and histone modifications for *FRZB* and their ploySINE. **c** Bar plot displays the population frequency of the polySINEs in the first exon of *RUNX3*. **d** Overviews of nine candidate genes under local adaptation. Bar charts indicate the population frequency of candidate polySINEs and the average TPM values of their corresponding candidate genes across tissues. **e** The scatter diagram displays the linkage disequilibrium between T-SNPs and polySINEs. The x-axis represents the chromosome, and the y-axis represents the r2 values.

Importantly, we found a fixed polySINE at the first exon of *RUNX3* in the Goettingen miniature pigs and MiniLEWE, which was also detected in Southern Chinese domestic pigs, particularly in the Luchuan pigs (**Fig. 7c**). The *RUNX3* gene was famous as a tumor suppressor in a human T-cell malignancy^102^ and was a key part of the TGF-**β** induced signaling pathway^103^. In addition, 30 PCGs showed the extreme *Fst* in the long-term laboratory-inbred Bama Xiang pigs^104^and were significantly enriched in the AMPK signaling pathway (Corrected *P-*values = 0.00295, Table S8), and especially the *SLC3A2* (upstream) and *SIRT1* (UTR3) genes had the polySINEs with a perfectly fixed frequency.

Nine of 75 candidate genes could be mapped to known QTX (Quantitative Trait Loci/Gene/Nucleotide) data that were potentially associated with the phenotypic traits^105^ (**Fig. 7d**). We observed that four PCGs related to laboratory-inbred Bama Xiang pigs were associated with fat content and body weight, which was consistent with the direction of selective breeding in laboratory^104^. Especially, the *LEP* gene was highly expressed in adipose tissue and can produce a hormone called leptin, which was involved in the regulation of appetite, fat storage, and body weight^106^. Overall, these findings demonstrate that polySINEs can serve as a valuable source for studying genomic ancestry and local adaptive evolution in pigs.

### Mapping young SINEs to the genetic associations of complex traits

To explore the association of polySINEs with complex traits, we first collected a total of 4,072 trait-associated SNPs (T-SNPs) from 79 published GWAS studies of 97 complex traits of economic value in pigs, including 18 reproduction, 22 production, 36 meat and carcass, six health, and two exterior traits (**Fig. S30**). As shown in **Fig. 7e**, we identified 127 trait-associated polySINEs (T-polySINEs) that were in linkage disequilibrium (LD, r^2^ > 0.3) with T-SNPs among 296 domestic pigs (109 Asian and 187 European domestic pigs). In particular, it was found that these T-polySINEs were more likely to be enriched in the TxFlnkWk (Weak transcribed at gene), indicating that they have the potential for gene regulation (**Fig. 8a**).

**Figure 8.**
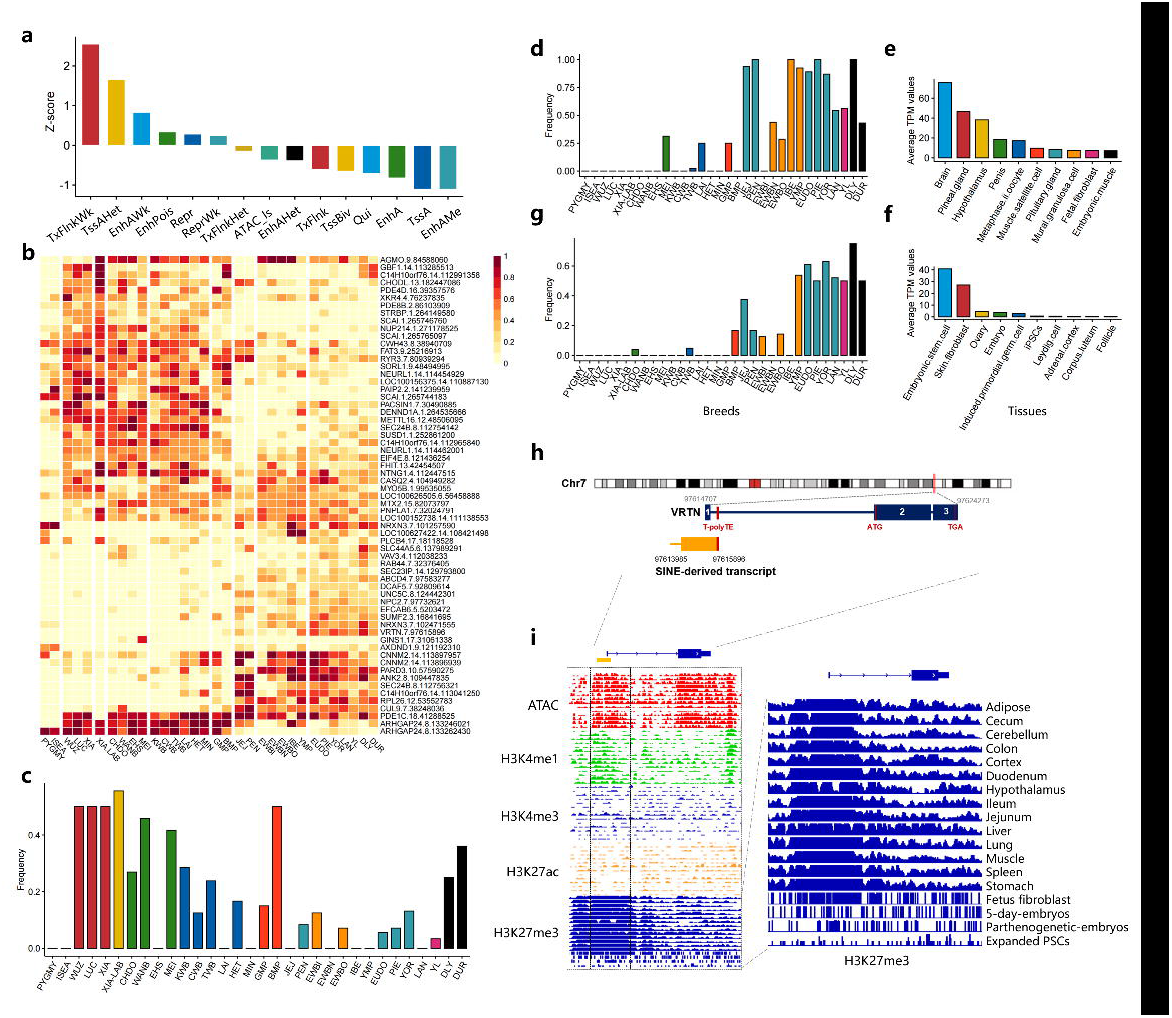
Mapping young SINEs to the complex traits. **a** Bar plot displays the enrichment of T-ploySINEs in different chromatin states. **b** Heatmap displays the frequency of T-polySINEs among different pig populations. The darker red color represents a higher population frequency for T-polySINEs. **c** Population frequency of T-polySINEs in *ELOVL3* gene among different pig populations. **d** Population frequency of T-polySINEs in *ANK2* gene among different pig populations. **e** The expression of the *ANK2* gene at the top 10 tissues sorted by gene expression. **f** The expression of the *VRTN* gene at the top 10 tissues sorted by gene expression. **g** Population frequency of T-polySINEs in *VRTN* gene among different pig populations. **h** The *VRTN* gene structure and the neighboring SINE-derived transcript. **i** Chromatin accessibility and histone modifications for the upstream of *VRTN* gene. H3K27me3 signals for the upstream of *VRTN* gene.

A total of 54 genes were affected by these T-polySINEs, which were associated with the intramuscular fat composition and teat number. The majority of them showed specific expression in certain tissues (Z-score > 2), particularly in the nervous system (plasmacytoid dendritic cells, choroid plexus, hypothalamus, and brain), reproductive system (testis, oviduct, and oocyte), and muscle satellite cells (**Fig. S31**, Table S9). Importantly, most of the intronic T-polySINEs exhibited breed-specific MAF between Chinese and Western pigs, which was in agreement with their differences in fatty acid content and teat number (**Fig. 8b**).

We identified a 320kb T-polySINE hotspots (chr14:112,965,840-113,285,513; r^2^ > 0.3), including six T-polySINEs and eight genes, which were significantly associated with intramuscular fat composition (**Fig. S32**). Among these hotspot genes, C14H10orf76 (*ARMH3*, r^2^ = 0.89) and *GBF1* (r^2^ = 0.86) have been reported to be essential for Golgi maintenance and secretion^107^. The *ELOVL3* gene was a strong candidate gene for fatty acid composition^108-109^. A low-frequency polySINE was in the intron region of *ELOVL3*, while multiple T-polySINEs within its upstream region of 15kb to 50kb. In particular, the T-polySINE (chr14:113,199,425) near 27 kb upstream was at high frequency in Chinese domestic pigs, especially Southern Chinese domestic pigs (**Fig. 8c**).

Furthermore, there was a high level of pairwise LD (*r*^2^ = 0.88) between the T-polySINE (chr8:109,447,835) in the intron of *ANK2* and the T-SNP that had a significant association with the fatty acid content of C14:0, C16:0, and C16:1n7 in backfat^110^. It was reported that ankyrin-B (AnkB) was a neuron-specific alternatively spliced variant of *ANK2* that was associated with obesity susceptibility in humans^111^. We found that the insertion of this T-polySINE was located in an LD block of 15kb (r^2^ > 0.5, chr8:109,439,023-109,454,866, **Fig. S33**) and almost fixed in the Western domestic pig populations (**Fig. 8d**). *ANK2* gene was observed to be ubiquitously expressed in pig bodies and highly enhanced in the nervous system (**Fig. 8e**). We noticed that two predicted SINE-derived transcripts (supported by Iso-seq reads) overlapped with exons of *ANK2* had a significant correlation with the expression of *ANK2* (Table S10). The expression of *ANK2* was significantly upregulated in cultivars with high-fat deposition compared with those with low-fat deposition, such as Songliao black pigs vs. Landrace pigs^112^, and fast-growing chickens vs. slow-growing chickens^113^.

In addition, we found a high LD (r^2^ = 0.75) between a T-SNP (chr7:97,606,621) and a T-polySINE (chr7:97,615,896) in the first intron of *VRTN* gene, which was significantly associated with teat number^114^. The *VRTN* was proposed as the most promising candidate gene to increase the number of thoracic vertebrae (ribs) in pigs^114^, which was highly and specifically expressed in embryonic stem cells, embryo, and ovary (**Fig. 8f**), suggesting that *VRTN* functions at the early embryonic stage of pig development. We found that this T-polySINE showed the obvious difference in frequency between Chinese indigenous breeds and commercial breeds (**Fig. 8g**). Especially, a novel transcript (length = 2,191bp) derived by this T-polySINE and covered the first exon of *VRTN*, which was significantly correlated with the expression of *VRTN* (**Fig. 8h**, *P-*values = 3.03×10^−201^, Table S10), and this transcript was supported by the RNA-seq exon coverage in NCBI annotation (**Fig. S34**). Of particular note, this region exhibited the open chromatin and enhancer signals (H3K4me1) while was facultatively repressed in most tissues (H3K27me3) (**Fig. 8i**). We clearly observed a significant decline in the repressed states at the stem cells and embryo-related tissues, which corresponded to the tissue-specific expression in *VRTN*, suggesting that this region was crucial for *VRTN*, and this T-polySINE was more likely to affect its expression.

## Discussion

### TEs are major, abundant, and polymorphic in the pig genome

In this study, we built a comprehensive atlas of TEs in the pig genome by using the newly developed PigTEDC pipeline, which combined the similarity-, structure-, and *de novo*-based methods. Our results demonstrated that nearly a third (947 MB) of the pig genome was made up of TEs, and the majority of them were non-LTR retrotransposons (SINE and LINE). SINE was shorter and more complete than LINE. Similar to our previous findings^53^, SINE, especially the PRE1-SS, PRE0-SS, and PRE1a families in SINE/tRNA, displayed the most recent ages and most polymorphic insertions. These polymorphic SINEs contributed nearly 90% of medium-length SVs among different assemblies, especially the L13 subfamilies, with 36.15% that were classified as the youngest SINEs in the pig genome.

### The influence of SINEs on transcriptional networks are associated with their ages

Gene regulatory network is influenced by genomic components, chromatin accessibility, histone modifications, DNA methylation, and *cis*-regulatory elements (e.g., TFBS, promoters, and enhancers). TEs associated with unique chromosome features can contribute to gene regulatory networks in a variety of the above ways. This is the first time, to our knowledge, to use large-scale multi-omics data to fully explore the relationships between TEs and chromosome features in the pig genome. Our findings showed that SINEs were significantly enriched in the A compartment, and the enrichment of SINE in open or closed chromatin regions was associated with their ages. For example, young SINE families were frequently enriched in close chromatin-like nucleosomes but highly depleted from open chromatin.

As expected, SINEs were highly depleted from all active chromatin tags, and more signals of constitutive heterochromatin tags (H3K9me3 peaks) were observed on SINE. The exception was H3K27me3, which was associated with facultative suppressor genes and cannot make permanent silence on SINEs. Almost all signals of histone modifications in SINE showed a tendency to decay as the age of TE increased, which was in line with the contribution of SINE to TFBS and the distribution of DNA methylation on SINE. However, there is still evidence that some young SINEs enrichment in weak functional regions, such as the youngest SINEs families, were highly enriched in the regions of weak active enhancer at hypothalamus tissue (Fold >1.5). We speculate that the relationship between SINE and its host genome is a combination of both “arms race” and “co-evolution,” depending on how the symbiosis turned out. In the former case of parasitism, the young SINEs were more likely to be treated as new invaders that were constitutively silenced by histone modifications and DNA methylation of the host genome (e.g., PIWI -piRNA pathway during the TE in testis), while the old SINEs were mutated and gained new regulatory potential, and thus tolerated or even co-opted by the pig genome. In the latter case of mutualism, there might be rare cases where the SINEs were positively selected by nature thereby help the host genome better adapt to the local environment in the long run.

### The non-coding RNAs derived from young SINEs may affect tissue-specific genes

The use of long-read isoform sequencing provided us a more complete characterization of full-length transcripts, which made it possible to identify the young SINE-derived transcripts. Meanwhile, the Iso-seq reads we used here were collected from about 40 pig tissues, which ensured the investigation of abundance and tissue specificity of young SINE-derived transcripts.

Our findings showed that the vast majority of young SINE-derived transcripts were non-coding RNAs that covered exons or felled in the introns. A total of 3,112 PCGs were found to be associated with young SINE-derived transcripts, and nearly 88% of them were enriched in co-expressed modules with high tissue specificity. The young SINEs that derived transcripts exhibited lower CG methylation levels and were more significant enrichment in open chromatin and histone modifications than whole young SINEs. Especially, some young SINEs exhibited strong and accordance tissue specificity in both transcript expression and epigenetic regulation. This was consistent with previous findings in other species^115-117^, suggesting that SINE insertions may be a crucial component of genes and regulate tissue-specific expression of their target genes.

### The detection of polySINE is significantly affected by detection tools and sequencing depth

PolySINEs belong to SVs that were more sensitive to sequencing depth than SNPs. In this study, to ensure the unbiased detection of polySINEs, we benchmarked the four polySINE detection tools under different sequencing depths. Our findings showed that MELT had robust performance in both *ref*+ and *ref*-detection (**Fig. S17-S18**). As expected, we found that the detected number of polySINEs increased significantly with the sequencing depth, especially from 5× to 10× that nearly doubled the average number of polySINEs (**Fig. S19**). Considering the sequencing depth in the current 838 publicly available whole-genome sequence datasets in pigs, we retained 374 individuals whose sequencing depth was greater than 10× and down-sampled their sequencing depth to ∼10× through a strategy of randomly removing reads (average mapped bases: 27.17 GB and average mapping rates: 99.43%). Finally, the PigTEP pipeline was developed to identify both polySINEs and SNPs in individuals simultaneously. This pipeline will help other researchers to explore the role of polySINEs in pig genomic study and breeding.

### TEs are non-negligible genetic markers for genetic diversity and complex traits

In pig genomic researches, the contribution of TEs to genetic diversity and complex traits was underestimated, even though SINEs can be more active when the pig is under selective pressure. The lack of a comprehensive map of polySINEs based on the large-scale re-sequencing dataset has limited our understanding of SINEs in pig population genetics.

Here, we genotyped and analyzed 211,067 polySINE loci in 374 individuals across 25 pig populations and found that polySINE loci were highly variable in polySINE allele frequencies among populations. The genetic relationships of samples based on polySINEs were consistent with previous studies based on SNP genotypes^118^, revealing that polySINE loci were informative in studying population genetics. Especially, we observed that ten major clusters representing 25 pig populations well corresponded to the geographic differentiation. These polySINEs with high pairwise *Fst* value were useful resources to understand local adaptation in domestic pigs. Our findings confirmed the results of previous studies (e.g., *IGFBP7*) and provided novel candidate genes that are potential to contribute to economically complex traits in pigs.

The genome-wide association studies (GWAS) based on SNPs have discovered thousands of QTLs of important economic traits in pigs, but most of these loci have not been functionally characterized. One possible reason is that what really affects phenotypic changes is not SNPs but SVs (MEIs) that were in linkage disequilibrium (LD) with them. In this study, 127 polySINEs were found to be LD with significant GWAS SNPs of complex traits, and nearly a third of them showed high tissue specificity in terms of expression. Importantly, a part of these polySINEs has been found to have the ability to derive novel functional transcripts (in H3K4me1 and H3K27me3), as the exon-covered transcript was found in the upstream of the *VRTN* gene. Future researches will be required to functionally validate whether and how these polySINEs affect complex traits by regulating their target genes in particular tissues (e.g., *VRTN* gene in embryonic stem cells).

## Acknowledgements

This work is financially supported by the Sanya Yazhou Bay Science and Technology City (202002007), the National Natural Science Foundation of China (31941007), and the AFRI grant numbers 2019-67015-29321 and 2021-67015-33409 from the USDA National Institute of Food and Agriculture (NIFA). We thank Sylvain Foissac from Université de Toulouse and other members associated with the FRAGENCODE project for their contributions to data of chromatin accessibility used in this manuscript.

## Author’s contribution

P.Z., Z.W., L.F., and G.L. conceived and designed the experiments. P.Z. designed analytical strategy and performed all analysis process. Y.G. assisted the DNA methylation analysis. Z.P. assisted the Histone modifications analysis. L.L. and X.L. assisted the bioinformatics analysis process. L.G., H.Z., X.H., and L.Q. collected and prepared for sequencing data. P.Z., G.L., L.F., Z.W., H.Z., and D.Y. wrote and revised the paper. All authors read and approved the final manuscript.

